# Decoding Spatial Attention in the Cocktail Party Problem Using Wearable Whole-head High-Density fNIRS

**DOI:** 10.64898/2026.05.06.722322

**Authors:** Sudan Duwadi, De’Ja Rogers, Alex D. Boyd, Laura B. Carlton, Yiwen Zhang, Anna Kawai Gaona, Aneesa Diya Pathiyaparambath, Ravin Chaudhury, Grace Magee, Bernhard B Zimmermann, Walker J O’Brien, Alexander von Lühmann, David A. Boas, Meryem A. Yücel, Kamal Sen

## Abstract

Spatial attention is critical for solving the cocktail party problem, a longstanding problem in neuroscience and artificial speech recognition. The ability to decode where humans are attending in a cocktail party like scene would empower applications in brain computer interfaces and assistive devices such as hearing aids. Here we demonstrate that, in an overt attention task, the attended spatial location can be decoded robustly from single trial hemodynamic responses, using a wearable whole head high density fNIRS system. We also identify critical brain regions that make the highest contribution to decoding accuracy. Specifically, we find that decoding based on a small fraction of channels within the left and right inferior parietal lobule (IPL), achieve maximal decoding accuracy comparable to all channels. These results open the way for the design of novel BCIs and assistive devices integrated with fNIRS, that can be steered by spatial attention.

## Introduction

The cocktail party problem (CPP), the problem of following a target speaker in a cluttered scene amongst other competing speakers (Cherry, 1953; McDermott, 2009), is a challenging problem for many humans, e.g., those with hearing loss (Arlinger, 2003; Podury et al., 2023), ADHD (Fu et al., 2022; Mihali et al., 2018) and autism (Marco et al., 2011). Normal hearing listeners can solve this problem with relative ease, indicating the existence of a solution in the brain. However, human brain regions that enable the solution remain poorly understood.

Spatial attention is thought to play a critical role in processing and analyzing complex cluttered scenes with competing audiovisual objects (Shinn-Cunningham, 2008), as in the CPP. For example, at a crowded gathering, a listener can look at and focus their attention on a target speaker at a spatial location to follow what they are saying. Relatively little remains known about human brain networks underlying spatial attention in the CPP.

Many studies have employed purely visual or auditory paradigms to investigate spatial attention. However, in complex everyday scenes, visual, auditory, and multimodal regions work in concert to select and process audiovisual objects. Indeed, a previous study found an effect of gaze direction on improving cocktail party listening (Best et al., 2023). In everyday scenes, we naturally orient towards an object by moving our eyes and then selectively process that target (overt attention). Here, we investigate the CPP using audiovisual stimuli, i.e., movies of competing speakers from different spatial locations, in an overt attention task. We also compare this with the processing of sounds of competing speakers, when the subject attends to a spatial location without visually orienting and looking at that location (covert attention).

Previous studies in humans have investigated the cocktail party problem using various methods, e.g., ECoG, EEG and MEG. fNIRS offers a number of advantages relative to other methods for investigating cortical networks in humans (Boas et al., 2014; Ehlis et al., 2014; Yucel et al., 2017). First, fNIRS is non-invasive and allows for studies in broader human populations compared to ECoG. Second, fNIRS is portable allowing for experiments in more naturalistic, ecologically valid settings compared to fMRI and MEG (Pinti et al., 2018; von Lühmann et al., 2021). Third, fNIRS has better spatial resolution compared to EEG, providing the potential to reveal specific cortical regions. A previous study provided a preliminary fNIRS dataset in a cocktail party-like setting (Ning et al., 2024). However, this study did not employ a high density system or monitor eye movements and did not identify specific cortical regions important for spatial attention. Here, we investigate the cortical representation of spatial attention in the cocktail party problem using a wearable whole-head high-density fNIRS system. We then decode the locus of spatial attention from the measured cortical responses using classification techniques. Such a system has potential for applications in brain computer interfaces for “steering” assistive devices, e.g., hearing aids, to the attended location to assist users with communication in noisy multi-speaker scenes. Additionally, such a system could enable attention-guided resource allocation for audio-visual rendering in AR/VR environments. Our results provide a glimpse into the human brain engaged in solving the cocktail party problem in more naturalistic conditions and reveal specific cortical regions that make an important contribution to decoding attended spatial location using fNIRS.

## Results

### Overt Attention Task

Our experimental design mimics an ecologically valid cocktail party scenario as shown in figure 1. In the overt scenario, three second audiovisual movie clips are presented simultaneously at 30 degrees to the left and right of the subject. Prior to each clip, a 2-second spatialized white noise cue is paired with a white crosshair on the corresponding screen, guiding subjects on which direction to focus, with eye movements allowed. A total of 26 participants were included in the main experimental analysis. An additional control experiment was performed in 8 returning participants (refer to the methods for inclusion and exclusion criteria).

**Figure 1:**
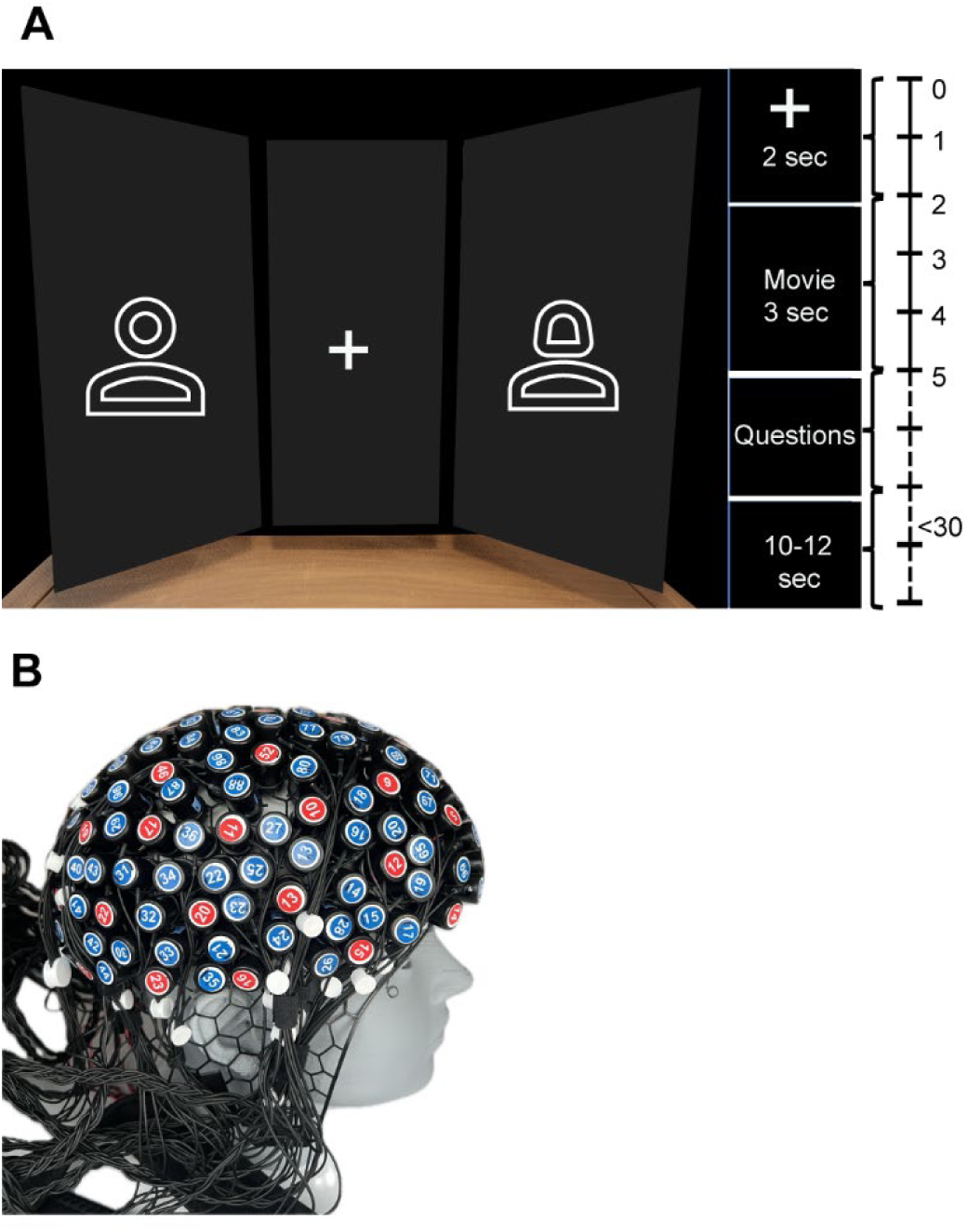
A) 3D rendered illustration of setup with a timeline of trial progression. The left side shows the schematic representation of experimental setup from subject’s point of view during one of the example trial blocks for overt conditions. The right side shows the timeline of a single trial progression. B) Picture of ninjaNIRS22 system on a mannequin head model for illustration.

#### Hemodynamic Evoked Response

We analyzed the evoked responses for all channels spanning the whole surface of the head to characterize cortical activity patterns. The processing pipeline (figure 10, methods section) was used to obtain the group average results shown in figure 2. All of the processing was done using combinations of custom written pipelines using Cedalion functions (Middell et al., 2026) and Homer 3 in MATLAB (Huppert et al., 2009) (analysis code available with link attached in Data availability section).

**Figure 2:**
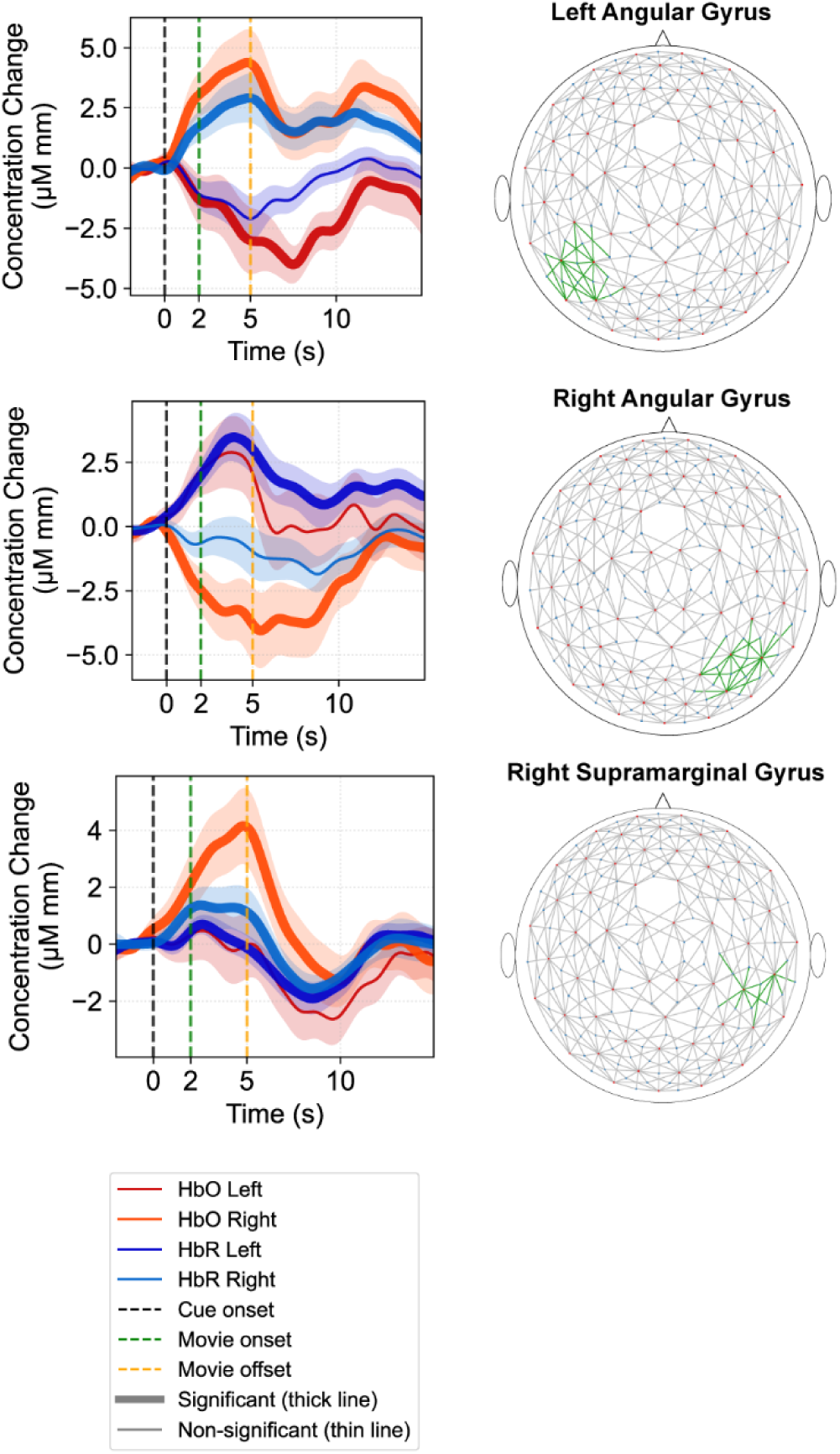
ROI average hemodynamic response functions on a group level. Only a selected few ROIs with statistically significant evoked responses are shown. Statistical significance was determined using t-test on the 2 seconds baseline (t=-2 to t=0) segment compared to the 2 seconds window starting cue (t=n to t=n+2, starting t=0) for both oxy and deoxy hemoglobin concentration. If any of the segments were statistically significant (p<0.05 after Bonferroni correction for multiple comparisons), the corresponding evoked response was marked as a statistically significant evoked response and indicated by bold lines. Ribbons represent standard error of the mean at group level.

Our results show statistically significant hemodynamic evoked responses in various regions of the brain during active engagement in the cocktail party problem. Selected examples are shown in figure 2 (see supplementary figures 2 and 3 for group level evoked responses of all ROIs in overt task). The accompanying 2D spatial plots show the spatial location of the corresponding ROI (colored in green).

##### Classification results

We decoded attended spatial location based on hemodynamic evoked responses using binary classification, where the two classes were ‘attended left’ and ‘attended right’ conditions. We used a random forest classifier with 10-repeat 5-fold cross validation. We first selected the top 20 channels via d-prime ranking, then applied Principal Component Analysis (PCA), and used max amplitude and slope of PC time-courses in moving 1-second windows (with 0.5 seconds step) as features (refer to methods section for more details). Mean classification accuracy (in percentage) with 95% confidence interval (figure 3) and latency (in seconds) to lower bound of 95% confidence interval (CI) to the classification peak are reported. An example of classification accuracy vs. time for one of the top five subjects is shown in figure 3.

**Figure 3:**
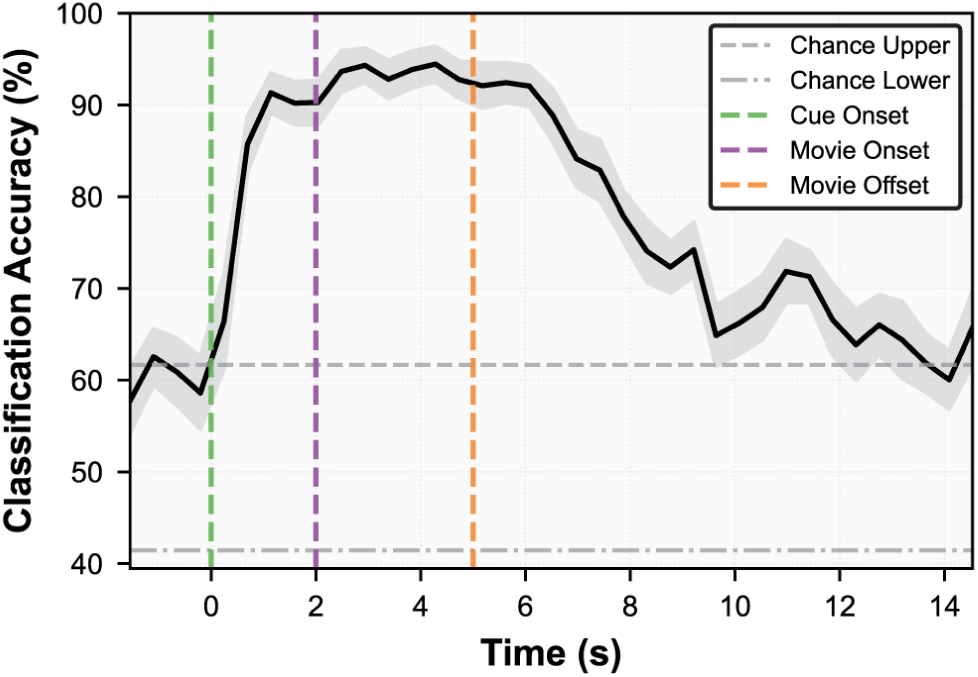
Example classification accuracy of an example subject in overt condition as a function of time. All channels spanning the whole head coverage were considered for this portion of analysis. Black line represents mean classification accuracy across 10 repetitions in 5-fold cross validation, and gray ribbon represents the standard error of the mean across the folds and the repetitions. Horizontal lines in gray represent upper and lower 95% confidence intervals for chance level (determined by permutation test). Vertical lines in green, purple, and orange represent cue onset, stimulus onset, and stimulus offset respectively.

###### All channels under consideration

Across all 26 subjects, mean classification accuracy was 76.5% (range: 52.3% to 100%), with 21 of 26 subjects exceeding the upper bound of the permutation-derived chance interval (62.3%; Figure 4A). Among the subjects performing above chance, the mean accuracy curve first rose above the lower bound of the 95% confidence interval of peak accuracy at a mean latency of 2.5 seconds (figure 4B).

**Figure 4:**
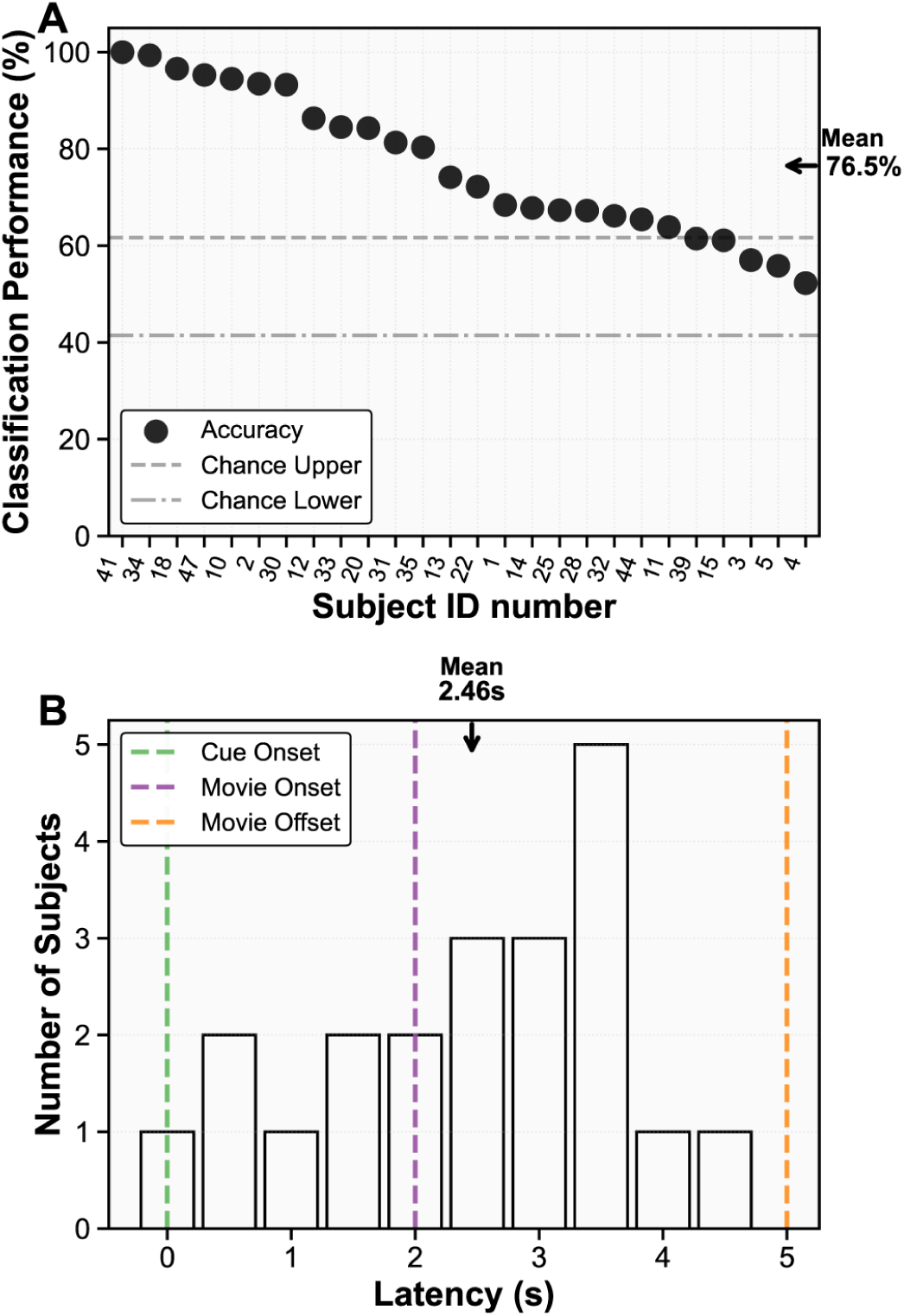
A) Mean from 10 repetition of 5-fold cross validated classification accuracy (expressed in percentage) is shown. Two horizontal gray lines represent upper and lower confidence interval for chance level after permutation test. B) Latency, defined as the earliest time the mean accuracy first crossed the lower bound of the peak’s 95% confidence interval (expressed in seconds), is shown as histogram. Latency values corresponding to the accuracy that were below chance level were excluded. All channels spanning the whole head coverage were considered for this portion of analysis.

#### ROIs contributing to the Classification

We quantified the contribution of each channel and each ROI to the classification performance in figure 5 (see methods). Right angular gyrus shows the highest mean contribution of 15% followed by left angular gyrus with mean contribution of 12% respectively, with both contributing in 100% of the subjects on group level (color coded as subject frequency in figure 5B). Other top ROIs included left visuomotor cortex, right premotor and supplementary motor cortex and right superior temporal gyrus.

**Figure 5:**
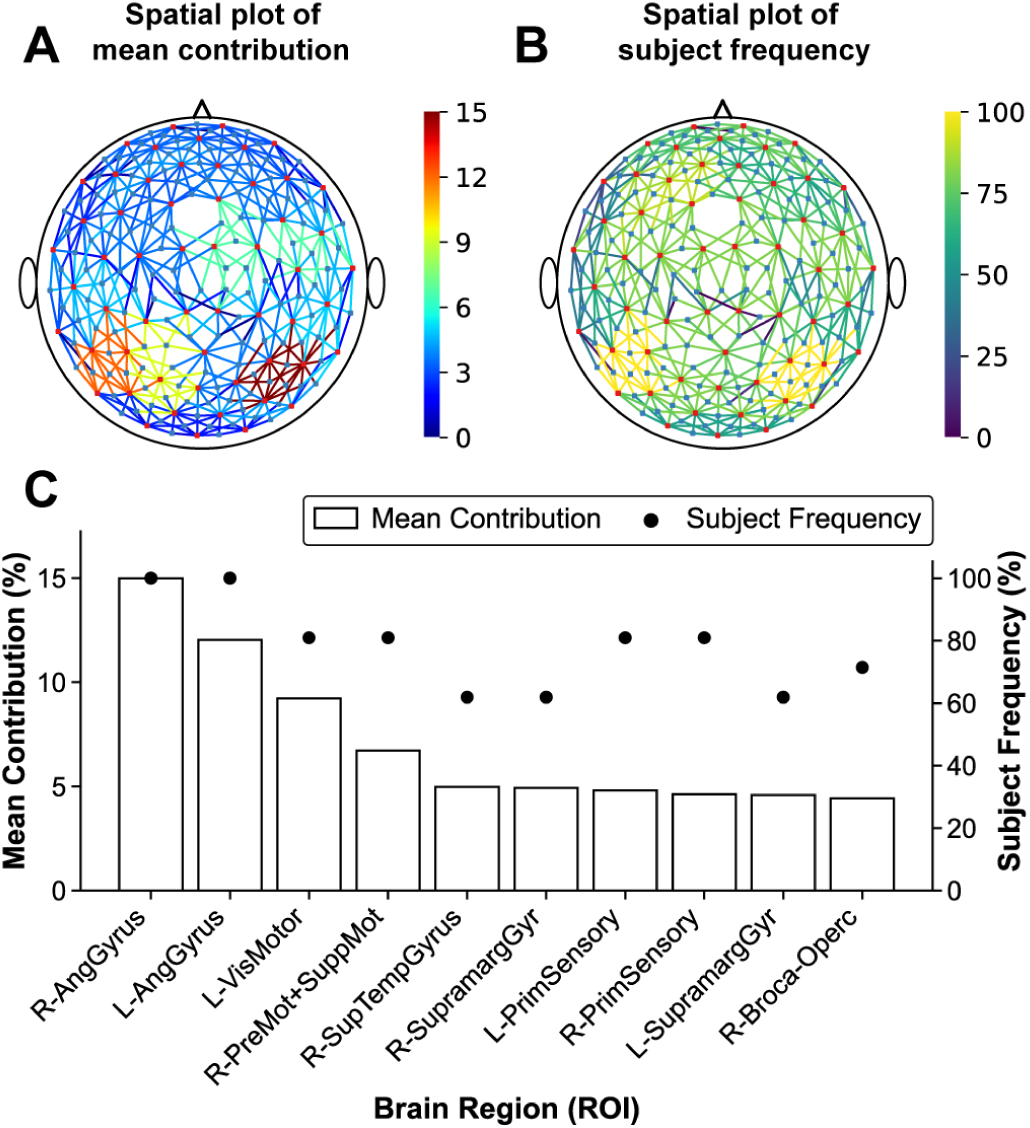
Mean contribution of ROIs at the group level. Principal component-based contribution is shown for each ROI at group level by aggregating importance from individual subjects’ ROIs (see methods for more details). A) Spatial 2D plot of mean contribution of ROIs. B) Subject frequency is shown in 2D spatial plot where max is 100% (given ROI appearing in all subjects) and minimum is zero (given ROI not appearing in any subjects). C) Top ROIs shown as bars, ordered by mean contribution on the left axis (only top 10 shown here). Subject frequency is shown as scattered dots on the right axis for corresponding ROIs.

##### Contribution of auditory component

Since our stimuli were audiovisual, we were interested in dissecting the contribution of the auditory component to evoked responses and classification performance. To assess this, we further analyzed the ROIs that made a relatively strong contribution to decoding performance (top 5 ROIs). We compared the evoked responses in the overt attention condition to a covert auditory condition (see supplementary figures 4 and 5 for group level evoked responses of all ROIs in covert task), where subjects listened covertly to audio only stimuli presented at the same spatial locations as the overt condition (left vs. right screens). In this condition, the subjects did not visually orient to the attended location but maintained fixation on the center screen. This condition eliminated both the visual stimuli and the visual orientation components in the overt condition, allowing us to assess the auditory contribution to evoked responses and classification performance. We looked at peak responses in top 5 ROIs on a group level to see if the responses were similar in the covert vs. overt conditions (figure 6). Figure 6 shows that most points in the scatterplots lie close to the diagonal, indicating comparable evoked responses in the covert auditory and the overt conditions (with the possible exception of left angular gyrus). Thus, the auditory component in stimuli drove evoked responses that are comparable to the overt condition. However, we found that classification performance (between left vs. right) in the covert condition was significantly lower than the overt condition (with mean performance = 55.9%, paired t-test, p < 0.001; see supplementary figures 7 and 9). These results indicate that, in the overt condition, visual components (visual stimuli and/or visual orientation) make a dominant contribution to classification performance.

**Figure 6:**
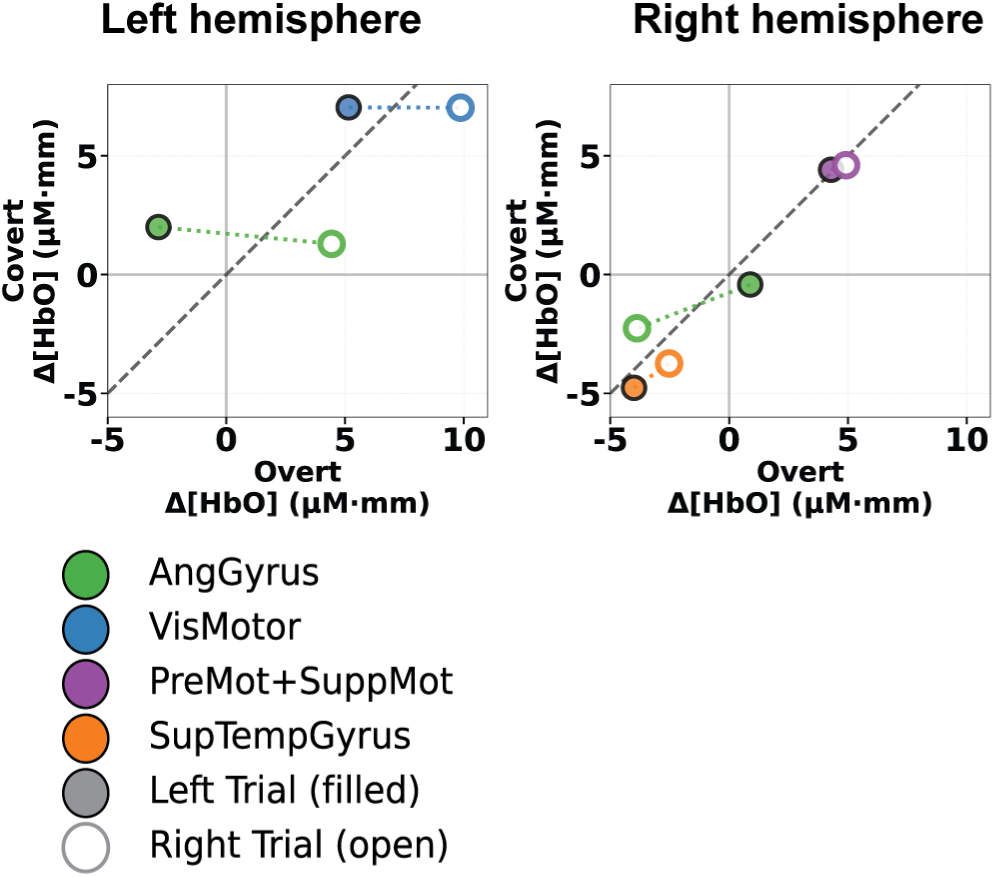
Scatter plots representing group-level peak hemodynamic amplitudes for the top five contributing ROIs. Each point represents the ROI-wise peak amplitude averaged across subjects for the two conditions plotted on the axes. Values closer to the diagonal indicate similar responses between conditions, whereas deviations from the diagonal indicate condition-specific modulation. Open circles denote attend-right trials and filled circles denote attend-left trials. Color-coded dotted lines connect points corresponding to the same ROI to aid visual correspondence across conditions.

##### Classification based on IPL channels only

Our ROI analysis indicated that both the left and right angular gyrus made a strong contribution to classification performance. We next assessed classification performance based only on these top two ROIs. The same processing and classification pipeline was applied as previously, except that only channels from a subset of the IPL region were included. An example of classification accuracy vs. time (for the same subject as shown in figure 3) is shown in figure 7, showing similar classification accuracy compared to all channels. At the group level, classification performance using IPL only channels vs. all channels, was not significantly different (paired t-test, p=0.29, supplementary figure 8).

**Figure 7:**
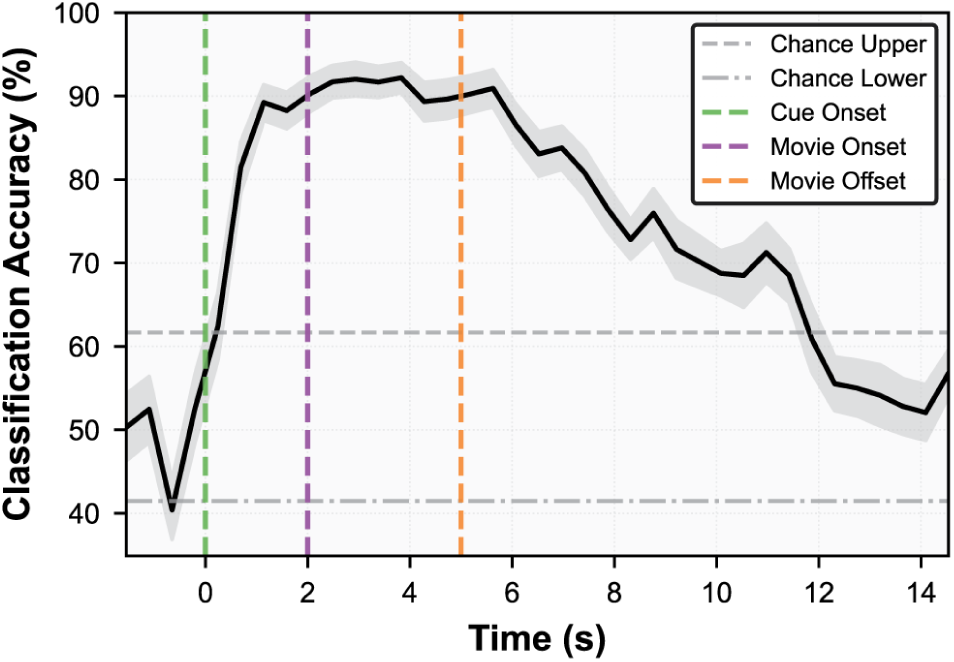
Example classification accuracy (of the same subject shown in figure 3) in overt condition as a function of time. Only channels covering left and right angular gyrus (subset of IPL) were considered for this portion of analysis. Black line represents mean classification accuracy across 10 repetitions in 5-fold cross validation, and gray ribbon represents the standard error of the mean across the folds and the repetitions. Horizontal lines in gray represent upper and lower 95% confidence intervals for chance level (determined by permutation test). Vertical lines in green, purple, and orange represent cue onset, stimulus onset, and stimulus offset respectively.

## Discussion

In this study, we attempted to decode the attended spatial location in a cocktail party like scene using whole head high density fNIRS measurements and classification methods. In the overt attention condition, we obtained robust classification performance based on single trials in most subjects. These results demonstrate the viability of a fNIRS based system for decoding attended location in a scene and opens the way to potential applications in brain computer interfaces for steering assistive devices to the attended location. We found that the average latency to peak classification accuracy in the overt condition was ∼2.5 seconds, which is within behaviorally relevant timescales in natural conversations.

Our results also shed light on specific brain regions important for decoding attended location, which is difficult to obtain using EEG which has poor spatial resolution compared to fNIRS. Specifically, we found that the classification accuracy obtained using only the channels from the inferior parietal lobule (IPL) region, was not significantly different from all channel classification accuracy. This result is consistent with the attention literature suggesting an important role of IPL as part of the fronto-parietal network in attention. Our results demonstrate that this area is accessible to fNIRS and plays a critical role in decoding attended location. It also opens the possibility of designing compact fNIRS devices specifically targeting the IPL for decoding spatial attention.

We observed significant variation in classification accuracy across subjects. Classification accuracy was significantly correlated with within-subject variability of the single-trial HbO response, quantified as the coefficient of variation (CV = σ/|μ|) computed across trials per channel, both when averaged across all channels and when restricted to channels overlying the Angular Gyrus (Supplementary Figure 6). Subjects whose single-trial responses were more variable relative to their mean amplitude tended to achieve lower classification accuracy. Thus, lower classification performance in some subjects may have resulted from less consistent trial-to-trial activation in key regions for decoding such as the IPL. Optimizing the collection of signals in key regions, in combination with individual subject scans, may further improve decoding performance in future studies.

### Key role of IPL in decoding performance

This study revealed key contributing brain regions to decoding performance, which would be challenging in other modalities such as EEG. The top ROIs included Brodmann regions of left and right angular gyrus (AG), visuomotor cortex / Superior Parietal Lobule (SPL), premotor and supplementary motor cortex, superior temporal gyrus (STG), and supramarginal gyrus (SMG). However, given that angular gyrus is a relatively small region, and our estimate of the uncertainty of ROI localization, our conservative estimate is that these channels were within the inferior parietal lobule (IPL, defined as AG plus SMG). The IPL is a key node in both the orientation network and the executive network underlying attention (Petersen & Posner, 2012), has been implicated in both sensory-motor integration and sound location monitoring during auditory spatial tasks (Alain et al., 2008), and carries direction-specific information during auditory spatial attention (Kong et al., 2014). The SPL (BA 7), a core node of the dorsal attention network, is known to exhibit transient activation time-locked to voluntary auditory attention shifts in both spatial and non-spatial domains (Shomstein & Yantis, 2006), with a spatial processing component observed during cocktail party listening (Hill & Miller, 2010). The premotor cortex has been implicated in the perception of acoustically degraded speech stimuli (Peelle, 2018); the stimuli in our experiments were also acoustically challenging due to competing stimuli from another movie. Thus, the top ROIs contributing to overt decoding performance identified in this study are consistent with their roles established in the attention literature. This study demonstrates that these regions also contribute in more naturalistic, ecologically valid settings (compared to fMRI, for example), and that these regions can be accessed with a wearable and portable fNIRS system, opening potential applications to BCI.

### Oculomotor component in IPL responses

In natural scenes, we visually orient by moving our eyes to attend to an object of interest. In such scenarios, both the oculomotor and attention systems are active and intimately linked. An interesting and challenging question relates to the relative contributions of these two systems under such conditions. Studies on attention have carefully considered whether these systems are independent, identical or overlapping (Corbetta, 1998; Corbetta & Shulman, 1998). Evidence suggests that attentional and oculomotor systems are tightly related and may be identical or overlapping, rather than fully independent (Corbetta, 1998). Our analysis comparing overt vs. covert classification revealed that visual stimuli and/or visual orientation makes a strong contribution to decoding performance. Given the leading contribution of IPL to decoding performance, we were specifically interested in whether eye-movement alone could account for decoding performance in IPL. To further decouple eye-movement signals from evoked responses, we designed an additional set of controls, where we introduced a longer delay between the visual orientation and the movie onset. This design allowed us to examine evoked responses long after eye movements were completed. Data from these additional control experiments revealed that responses in IPL are difficult to account for based on eye movement only (see supplementary methods, results, and supplementary figure 1). Rather such responses may represent attended location and/or eye position (which are identical in the overt condition). Such a representation may be similar to and arise from attentional gain fields found in neurons in parietal regions.

### Limitations and Future Directions

While we took steps towards a more naturalistic design for investigating the cocktail party problem by incorporating audiovisual stimuli and allowing for visual orientation using eye movements, additional factors arising in natural conditions remain to be explored. For example, future studies should investigate designs that allow for head movements which could occur under natural conditions.

Second, although the decoding of overt attention was robust, we did not obtain robust decoding for the covert attention condition. There are several possible explanations for this. In everyday natural scenes, we visually orient and attend to particular objects in the scene to analyze them in detail. In the overt condition, the visual orientation component and the visual component of stimuli may have generated a stronger signal, which in turn improves decoding. These components were absent in the covert condition. It is also possible that our fNIRS cap did not optimally target auditory cortical regions near the ear which could be sensitive to auditory only stimuli from different spatial locations. Future studies should investigate whether improved coverage of auditory cortical regions improves decoding performance, especially in the covert condition.

Third, this study used decoding based solely on fNIRS. Existing literature suggests that EEG-fNIRS sensor fusion techniques could potentially improve single trial decoding accuracy than fNIRS alone (Codina et al., 2025). We are currently exploring whether simultaneous measurement of fNIRS and EEG can further improve decoding performance in both overt and covert conditions.

Finally, this study was limited to binary classification between left vs. right. It will be interesting to expand this paradigm to a larger number of spatial locations and investigate the spatial resolution of decoding.

### Potential applications to BCI systems and assistive devices

Recently, we have proposed a brain inspired algorithm for sound segregation (BOSSA) (Chou et al., 2022). BOSSA significantly improves speech recognition in cocktail party-like settings for both listeners with normal hearing (Chou et al., 2022) and hearing loss (Boyd et al., 2025). BOSSA requires as input the spatial location a user wants to attend to. The decoded attended spatial location from fNIRS could be used to steer BOSSA to the attended location to better isolate the target sound from competing sounds. Similar steering of assistive devices, e.g., hearing aids, can be envisioned with the integration of fNIRS. The top ROIs for decoding attended location identified in this study, e.g., the left and right IPL, opens up the way for designing a more compact wearable fNIRS system which specifically targets these ROIs. Additionally, such a system could enable attention-guided resource allocation for audio-visual rendering in AR/VR environments and facilitate implicit and neuroadaptive brain–computer interfaces.

## Methods

### Participants and Procedures

The experiment was conducted in accordance with the Institutional Review Board of Boston University, ensuring ethical standards in conducting studies for human subjects. All the risks and benefits of participation were described to the subjects in advance and on the day of data collection. All the participants signed an informed consent form. Participants were screened for neurological or psychiatric disorder, history of hearing problems, cochlear implant usage, uncorrected vision problems, psychoactive substance usage, pacemaker or implantable defibrillator usage, and deep brain stimulator usage. Participants were stabilized using a chin rest and instructed to minimize movement throughout the task. All participants were compensated for their time.

Forty-three unique participants were consented and recruited for the main experiment. In addition, eight participants from the main cohort returned for the control experiment, yielding 51 attempted recording sessions in total. After exclusions due to hardware issues, participant dropout, lack of task comprehension, poor signal during calibration, and experimenter error, 34 usable recording sessions were included in the final analyses: 26 main-experiment participants and 8 control-experiment sessions from returning participants, as shown in Figure 8. The main analysis included 26 participants (20 male, 6 female; age range: 19–62 years). The control experiment included 8 returning participants (6 male, 2 female).

**Figure 8:**
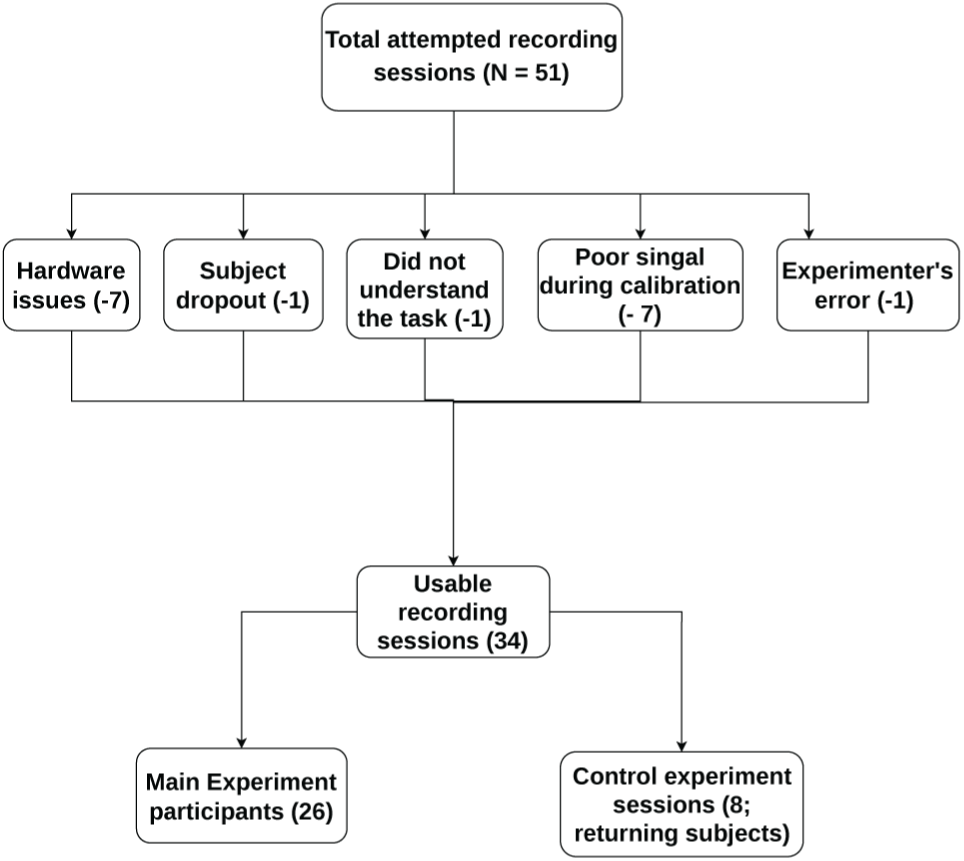
Flowchart of recording-session inclusion. A total of 51 recording sessions were attempted, including main experimental sessions and control sessions from returning participants. After exclusions, 34 usable recording sessions were included in the final analyses: 26 main-experiment participants and 8 control-experiment sessions from returning participants.

### Stimuli

The stimuli used in this study were extracted from AVSpeech, a publicly available audiovisual speech dataset (Ephrat et al., 2018). The videos were mined using a custom automated downloading script. Downloaded videos were automatically screened for presence of a single talker in the video, English as a spoken language, context of the language being not offensive, and sentences with at least 4 words that make semantic sense. Videos were further required to show the talker positioned approximately at the center of the frame, defined as cases in which the automatically detected face never entered the leftmost or rightmost third of the screen. All the custom scripts were written in Python with semantic and sentiment analysis done using OpenAI’s GPT-4 API and language detection (speech detection) using OpenAI’s whisper model. A total of 360 unique videos were shortlisted after manual inspection and verification of all the above-mentioned automated criteria. The audios from the shortlisted videos were spatialized - 30 to the left and +30 degrees to the right using HRTF filters based on KEMAR head model using MATLAB.

### Experimental Design

Our experimental design mimics an ecologically valid cocktail party scenario in both overt and covert contexts as shown in figure 1. In the overt scenario, 3-second audiovisual movie clips are presented simultaneously at 30 degrees to the left and right. Prior to each clip, a 2-second spatialized white noise cue is paired with a white crosshair on the corresponding screen, guiding subjects on which direction to focus, with eye movements allowed. In the covert scenario, subjects are exposed solely to spatialized audio from the same set of movies. Here, the 2-second spatialized white noise serves as the cue, directing their attention, while they maintain a gaze on a central screen displaying a static white crosshair on a light gray background. A total of 60 trials (30 trials for attend-left and 30 trials for attend-right conditions) for both overt and covert conditions were collected. Additionally, 20 trials (10 trials for look-left and 10 trials for look-right) were collected as eye movement control trials where the subjects move their eyes to fixate at the center of left or right screen in absence of any audio or video stimuli. At the end of each trial, the subject answered multiple choice questions by pressing 1, 2, or 3 on the numeric pad of a keyboard to answer audio related questions (What did the target speaker say?) and video related questions (Who was the target speaker?). Audio questions contained 3 options with a transcript of the audio played during the trial where one transcript is from the target speaker (one to be attended), another from the masker speaker (one to be ignored), and the last one from the pool of 360 short listed videos. The video question contained corresponding speaker’s face screenshots from the video that was played. In overt condition, subjects answered both audio and visual questions, but in covert condition, they only answered audio questions. For eye control tasks, the subjects move their eyes 30 degrees to the left and right with the same trial structure, totalling 5 seconds of fixation on the directed screen.

### Experiment Execution

The experiment was conducted using custom scripts written in Python using Psychopy (code available with link on code availability section). Upon the participants’ arrival, they filled out a demographics questionnaire and signed an informed consent form. A quiet room was used where the participants sat down in a chair in front of 3 monitor setup. The subjects were instructed to put their head on chin rest and instructed to not move their head or any body parts during the task period. The subjects were allowed to do 2-4 practice trials of each task to ensure task comprehension. fNIRS caps with appropriate sizes were fitted to their heads. Cap size was adjusted by changing the length of extender clips bridging the two hemispheres at the midline. Tobii pro (Tobii, stockholm, Sweden) eye tracking glasses 3 (first 16 subjects) or neon (pupil labs, Berlin, Germany) eye trackers (rest of the subjects) were put on the subjects to track their eye gaze on the monitors. Insert earphones (3M eartones, USA) with earplugs were placed in subjects’ ear to deliver the audio and reduce the ambient auditory noise. Bright overhead room lights were turned off and warm-tone halogen lamps were used to minimize ambient optical interference and improve fNIRS signal-to-noise ratio during optimization and recording. In addition, black shower cap was used to cover the entire fNIRS cap to reduce ambient light leakage into the fNIRS recording. Manual maneuvering of the hair was done to improve signal calibration. Subjective decisions were made by the experimenters until satisfactory signals (around 60% channels being of good quality) were achieved during the calibration stage.

#### fNIRS data acquisition

fNIRS data were collected using a continuous-wave, in-house–developed wearable whole-head high-density fNIRS system, NINJANIRS22 (O’Brien et al., 2024). The system operated at two wavelengths, 760 nm and 850 nm, with a sampling rate of 8.988 Hz.

#### Optode layout

A total of 56 sources and 144 detectors were used, forming a whole head montage with 567 channels per wavelength. The optode layout was designed in Atlasviewer software (Aasted et al., 2015). The source and detector optodes were mounted in a custom 3D-printed flexible fNIRS cap, referred to as ninjaCap (von Lühmann et al., 2024). Monte Carlo simulation was carried out in Atlasviewer to generate the photon migration sensitivity profile (figure 9).

**Figure 9:**
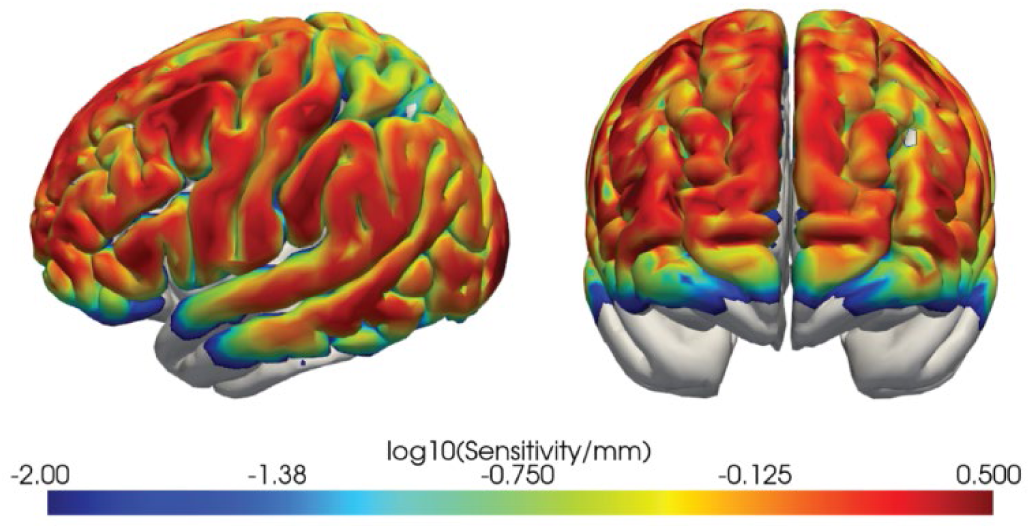
Sensitivity profile of the fNIRS optode layout. Monte Carlo Simulation of photons propagation was simulated using the MCX toolbox in Atlasviewer in Matlab.

#### Anatomical registration

In a separate subset of subjects with similar head sizes, we employed 3D photogrammetry to assess uncertainty in optode placement. Uncertainty was quantified by examining whether the post-registration centroid MNI coordinates of channels assigned to the inferior parietal lobule (IPL) remained within the anatomical IPL. Among the 10 subjects analyzed, only one subject in the left hemisphere and another subject in the right hemisphere showed centroid locations outside the IPL, indicating generally consistent IPL localization across subjects.

### fNIRS data preprocessing

fNIRS preprocessing was carried out in MATLAB programming language using Homer3 software package (Huppert et al., 2009) and in Python programming language using Cedalion toolbox (Middell et al., 2026). Raw intensity at both wavelengths was quality-screened for low signal and saturation; channels with amplitudes below 0.001 a.u. or above 0.84 a.u. were pruned. The remaining intensity data were converted to optical density (OD) and motion-corrected using Temporal Derivative Distribution Repair (TDDR) (Fishburn et al., 2019). The corrected OD was band-pass filtered between 0.01 and 0.5 Hz using a zero-phase Butterworth filter (order 4). Filtered OD was then converted to hemoglobin concentration changes using the modified Beer-Lambert law, without differential pathlength correction.

For group level average, the General Linear model (GLM) was used to estimate single trials by regressing out the systemic physiological noise using mean of all channels less than 20 mm source-detector separation as global signal regressor. The hemodynamic response was modeled using a sequence of Gaussian basis functions spaced 1 s apart with a standard deviation of 1 s, spanning −2 to +15 s relative to stimulus onset (Gagnon et al., 2011). The mean pre-stimulus baseline (−2 to 0 s) was subtracted from each trial. For each channel, all of the trials were averaged for each condition (left and right). For each subject, all of the channels were averaged within a defined region of interest (ROI). All of the ROIs were averaged across the subjects to get group average results.

### Eye Tracking data collection

Tobii pro 3 (Tobii, Stockholm, Sweden) eye tracking glasses (for the first 16 subjects) or neon (pupil labs, Berlin, Germany) eye trackers (remaining subjects) were used to collect eye tracking data. A calibration target provided with the eye tracking system was used to calibrate eye tracking before each recording session for Tobii glasses while no such manual calibration was done for Neon. Additional powered lenses were snapped on the glasses if subjects used prescription glasses and did not use contact lenses during their visit. Audio jack cable (3.5 mm) was used to connect the glasses controller to the fNIRS device to send trigger signals to the fNIRS device for syncing fNIRS and eye tracking data for Tobii. For the neon eye tracker, subjects were asked to nod their head at the beginning to align the accelerometer from the fNIRS cap and the neon.

### Tobii eye tracking data

Custom written MATLAB scripts were used to analyze eye tracking data. Horizontal gaze 2D data was used to determine gaze angles from the center of the screen. If the subjects moved their eyes more than 20 degrees tolerance to the left (-30 degrees) or to the right (+30 degrees), the trials were discarded. Example traces are shown in figure 11.

**Figure 10:**
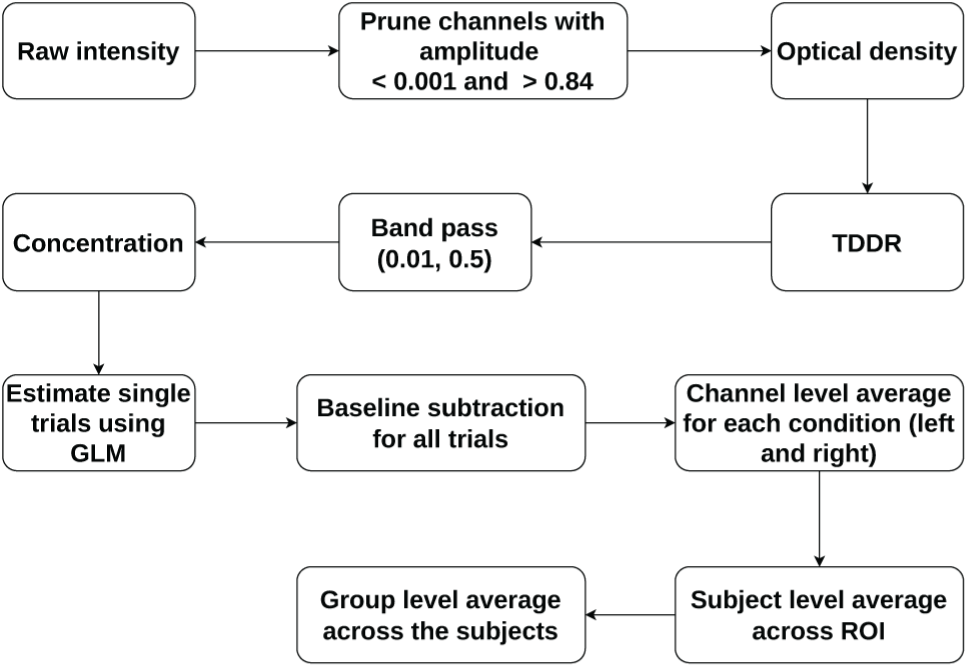
Preprocessing pipeline for fNIRS data processing to get Group level average across the subjects.

**Figure 11:**
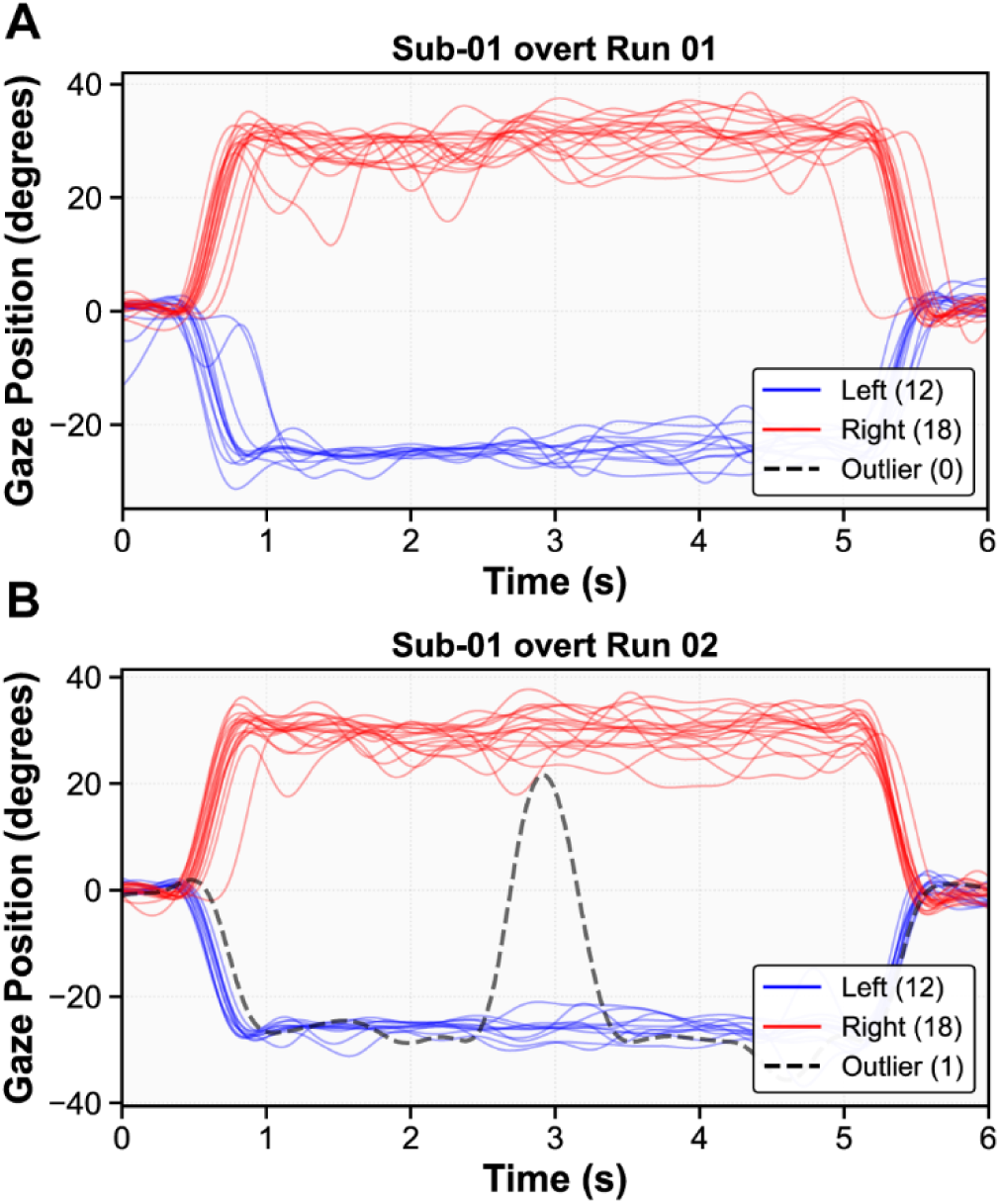
Eye tracking traces example for overt condition. Red color traces represent eye movement to the right and blue represents eye movement to the left. Black dotted line represents an outlier trial where the subject moved their eyes more than 20 degrees of tolerance. Panel A shows clean eye gaze traces without any outliers. Panel B shows one outlier marked in dotted line as an example.

### Eye Tracking data analysis

Publicly available software Glassesviewer (Niehorster et al., 2020) was used to parse the raw Tobii data into .mat readable files. The parse data contained sampling rate information (100 Hz), horizontal azimuth, and vertical elevation data among a few other pieces of information. For participants with data available from both eyes, left eye data was preferentially used, and right eye data was used when left eye data was not available. Only horizontal azimuth information was used for further analysis.

Raw gaze data was normalized by field-of-view (FOV) parameters (horizontal 95 degrees) and converted into gaze angles. Missing samples due to blinks or any other causes were replaced using linear interpolation. Signals were low pass filtered at 2 Hz using third order zero phase Butterworth filter.

Eye tracking data were temporally aligned to fNIRS data using TTL pulses recorded in auxiliary channels of the SNIRF files. The first detected pulse from the Tobii trigger signal at the onset of the Tobii data recording was used to compute the offset between stimulus onsets recorded in fNIRS data stream and Tobii data stream. For the neon eye tracker, subjects were asked to nod their head at the beginning to align the accelerometer from fNIRS cap and neon. Task condition specific (attend left and attend right) were matched to onset times for each run to extract relevant gaze data time course. For each trial, an epoch of -2 seconds to 15 seconds relative to the stimulus onset (t = 0 seconds) were used. For outlier detection, we thresholded with tolerance of 20 degrees for overt and control conditions where we expected -30 degrees and +30 degrees of eye movement. For covert condition, we thresholded with tolerance of 5 degrees. We restricted the time window of consideration to 2 to 4 seconds after the cue onset to avoid marking slower onset and offset of eye movements in response to the cue onset.

### Behavioral data analysis

Each multiple-choice question was marked correct or incorrect, separately for audio and video questions. Final correctness marking is based on answering both questions correctly for overt tasks and only audio questions for the covert task. Behavioral analysis was performed in MATLAB 2024b. Accuracy during the trial was recorded by the Psychopy custom script in real time during the experiment. CSV files with accuracy of response were used for post hoc analysis. Only correct trials were used for further analysis.

### Behavioral data

Each multiple-choice question was marked correct or incorrect, separately for audio and video questions. Final correctness marking is based on answering both questions correctly for overt tasks and only audio questions for the covert task. All the subjects scored more than 70% in the behavioral data metric, so no subjects were excluded based on behavioral response as shown in figure 12.

**Figure 12:**
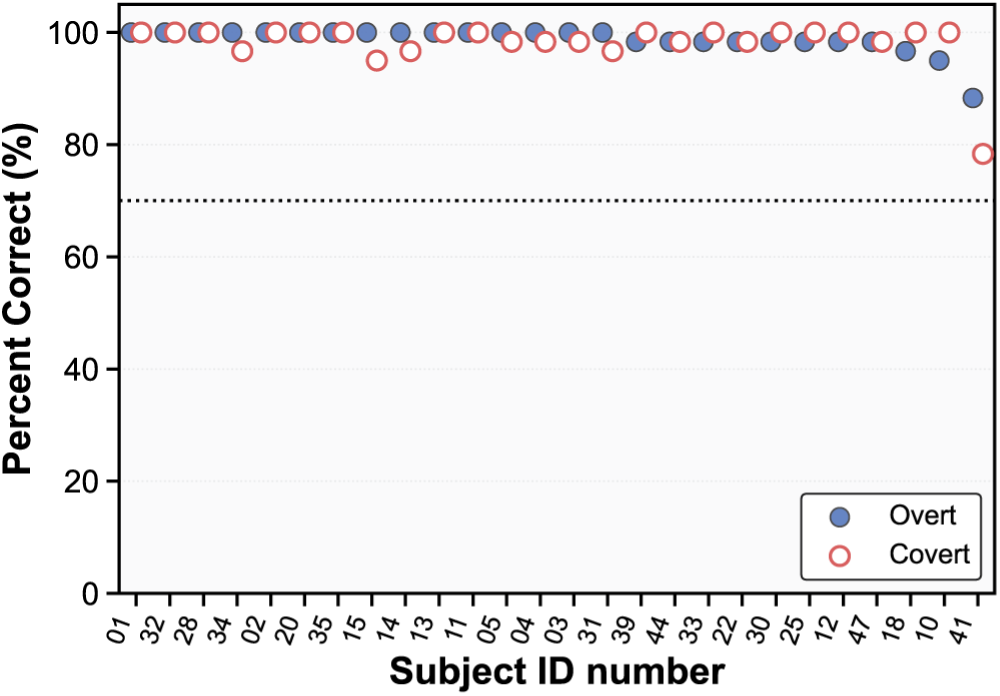
Behavioral accuracy across subjects for overt and covert attention tasks. Each dot represents behavioral accuracy, defined as correct responses to both auditory and visual questions in the overt task and correct auditory responses in the covert task. It is sorted by overt behavioral task performance. Black dotted horizontal line represents the 70% criteria to reject a subject based on behavioral correctness.

### Group level Average

Group level average was calculated by averaging all single trials across the channel space and across the subjects in each defined region of interest (ROI). Standard error of the mean was calculated across the subjects to report on hemodynamic response functions (HRF).

### Statistical analysis for evoked responses

Statistical significance was determined using a t-test on the 2 seconds baseline (t=-2 to t=0) segment compared to the 2 seconds window starting cue (t=n to t=n+2, starting t=0) for both oxy and deoxy hemoglobin concentration within each condition (e.g., left HbO). If any of the segments were statistically significant (p<0.05 after Bonferroni correction), the corresponding evoked response was marked as a statistically significant evoked response.

### Classification scheme

To avoid information leakage, a cross-validation scheme was employed where OD data was partitioned into training and testing datasets. The training dataset was filtered between 0.01 Hz and 0.5 Hz and converted into concentration. GLM was performed with global signal regression on only the training set. Estimated global signal regressor as systemic physiological contributor was used to regress out physiological noise from the training set on a single trial level. Identical to the group-level analysis, the HRF was modeled using a sequence of Gaussian basis functions spaced 1 s apart with a standard deviation of 1 s, spanning −2 to +15 s relative to stimulus onset (Gagnon et al., 2011). Single trials on the test dataset were estimated by subtracting global signal regressor weighted trials from the original single test trials (Ning et al., 2024). Baseline (-2 to 0 s) was subtracted from each individual trial.

To rank the features, the training set was used to rank the feature where d-prime (d’) is calculated as the absolute difference of the class means divided by the weighted sum of standard deviation (pooled standard deviation). Time windows of 2 to 10 seconds post cue onset were used to calculate d’ to avoid initial onset spike and late hemodynamic responses. d’ time course was further smoothed with 9 point moving average and thresholded by a hold criterion (sustained elevation relative to pre-window, 0 to 2 seconds post cue onset). Top 20 sets of channels were used based on top 20 highest d’ composite measures where d’ composite is calculated as 0.7*peak d’ + 0.3*area under the curve. Peak d’ captures maximum separation between classes and area under the curve captures sustained hold of the separation to avoid noisy d’. This method was used on each moving window of 1 second with stride size of 0.5 s for each dataset, thereby having different top 20 sets of features for each moving window across time. The features were passed to Random Forest (RF) binary classifier to classify single trials into left vs attend right classes. Mean accuracy on testing set over all the folds and repetition was used as the final classification accuracy for each moving window. A flowchart of classification pipeline is shown in figure 13.

**Figure 13:**
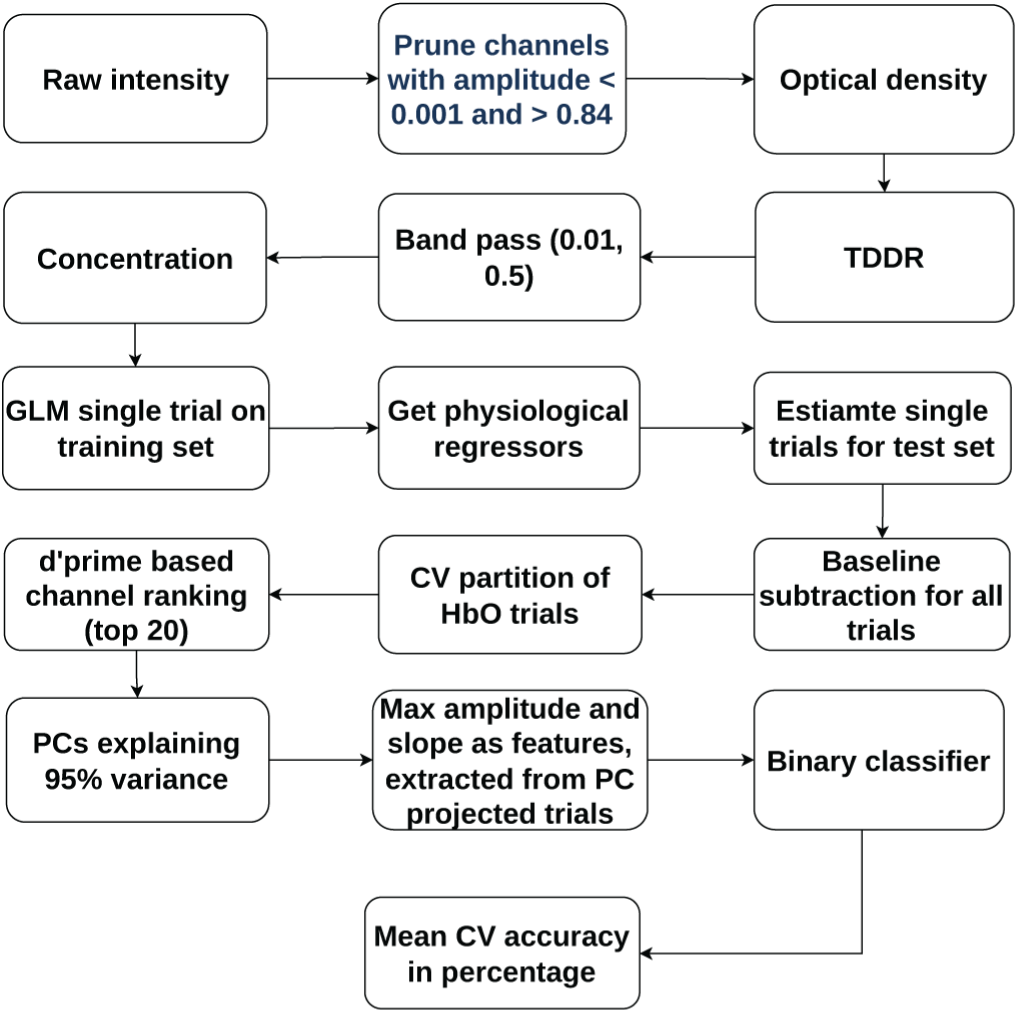
Flowchart for classification pipeline scheme

### Permutation test

Due to the small number of data points in the dataset (n = 60 trials total), the true chance level is not at 50% (Combrisson & Jerbi, 2015). Permutation test was carried out by shuffling the labels 1000 times (with 5-fold cross validation) to estimate the true chance level. Lower bound was estimated to be 40.7% and upper bound was estimated to be at 62.3% in one tailed test. Accuracy above 62.3% was marked as statistically above chance level accuracy.

### Feature Importance

#### Principal Component importance score

Within each outer fold, after baseline subtraction of each trial’s pre-cue mean, the selected channels (selected using d’ method) in training data were standardized by z-scoring column-wise across all training time samples. PCA was then fitted on the standardized training matrix, and the same standardization and PCA projection were applied to the test trials. Max amplitude and slope features were computed from PC time courses, which were obtained by projecting standardized (training-z-scored) channel data into PCA space. Standardization and PCA training were done only on training trials. Trained parameters (means, variances, component axes) were then applied to both training and test data. Channel importance was defined as the sum of squared loadings weighted by each component’s explained-variance ratio:

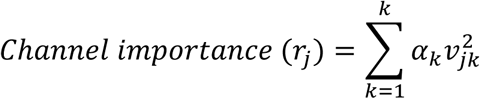

where *v*_*jk*_ is the *j*^*th*^ element of the unit-norm eigenvector for component k and *α*_*k*_ is that component’s explained-variance ratio.

For each subject and condition, we quantified the PC based contribution within each outer fold of cross validation (refer to methods for PC contribution calculation). Channel contributions were mapped to Brodmann ROIs and summed within each fold and averaged across the folds to get each subject’s mean ROI contribution. At group level, we computed the mean of the subject level means across the subjects in whom the ROI received nonzero contribution. In addition, we calculated ‘subject frequency’ as the fraction of subjects in which the ROI was present. Figure 5 ranks the top 10 ROIs by mean contribution. Bar height indicates group-level mean contribution, and dots indicate subject frequency, highlighting regions that are both strongly weighted and consistently observed across participants.

### AI Use Transparency

Large language models (Claude, Anthropic; ChatGPT, OpenAI) were used for grammar refinement of some manuscript sections and to assist with Python code development for the analysis pipeline. The authors reviewed and verified all AI-assisted outputs and take full responsibility for the manuscript content and associated code. The GitHub README further reiterates this, including docstrings on scripts where AI was used in any capacity.

## Data availability

Data are available on OpenNeuro.

## Code availability

Experimental setup with stimuli delivery scripts are available on [GitHub experiment](https://github.com/duwadisudan/cocktail-party-stimuli).

Data processing and analysis scripts are available on [GitHub processin](https://github.com/duwadisudan/wholehead-cocktail-party-fnirs/tree/v0.1-biorxiv)g.

Analysis provenance was tracked using whichscript, an open-source provenance tracking tool, available on [whichscript GitHub repository](https://github.com/duwadisudan/whichscript).

## Acknowledgements

We would like to thank Virginia Best and David Somers for advice on experimental design; Isaac Boyd and Shannon Kelley for code reviews; Sreekanth Kura for technical help and troubleshooting; Sharayu Shrinivas for assistance with the initial eye tracking code; Parya Farzam for administrative help; and Liam Caramello for help with the illustration in figure 1a.

## Author Contributions

**Sudan Duwadi**: Conceptualization, Data curation, Formal analysis, Investigation, Methodology, Project administration, Software, Validation, Visualization, Writing – original draft, Writing – review & editing.

**De’Ja Rogers**: Data collection, Preliminary analysis.

**Alex D. Boyd**: Experimental design, Investigation (experimental setup), Data collection, Eye tracking data analysis

**Laura Carlton**: Software (processing scripts), Data collection, Preliminary analysis.

**Yiwen Zhang**: Data collection, Preliminary analysis.

**Anna Kawai Gaona**: Data collection, Preliminary analysis.

**Aneesa Diya Pathiyaparambath**: Subject recruitment, Data collection, Preliminary analysis.

**Ravin Chaudhury**: Data collection, eye tracking data analysis

**Grace Magee:** Experimental setup and Data collection

**Bernhard Zimmermann**: Hardware development, Investigation (experimental setup).

**W. Joe O’Brien**: Hardware development, Investigation (experimental setup).

**Alexander von Lühmann**: Methodology (data processing and classification), Writing – review & editing.

**David A. Boas**: Conceptualization, Supervision, Funding acquisition, Writing – review & editing.

**Meryem A. Yücel**: Conceptualization, Supervision, Funding acquisition, Writing – review & editing.

**Kamal Sen**: Conceptualization, Supervision, Funding acquisition, Writing – review & editing.

## Funding

NSF 2319321

## Declaration of Competing Interest

K.S., S.D., M.A.Y., D.A.B., and Boston University have filed a provisional patent application related to the methods described in this manuscript (BU-2025-067, US Application No.: 63/916,093, "Device and System for Decoding Attended Spatial Location"). The remaining authors declare no competing interests.

## Notes

https://openneuro.org/datasets/ds007738/

https://github.com/duwadisudan/wholehead-cocktail-party-fnirs/tree/v0.1-biorxiv

### Summary of Updates

Added co-author; fixed typos; revised manuscript text, references, figure callouts, methods wording, and figure captions.

