## Supplementary for "Decoding Spatial Attention in the Cocktail Party Problem Using Wearable Whole-head High-Density fNIRS"

### Title

*Supplementary methods for additional controls*

To further disentangle the contribution of eye movement to evoked responses in the overt condition in IPL, we ran a longer control task where we invited 8 subjects from the same pool of 26 subjects we used for our final analysis. The experiment design largely remained the same as described in the main text body (see experimental design and execution section) except that the subjects were asked to maintain the eye fixation on the plus sign on the monitor of the cued spatial location for 15-17 seconds before the movie onset. In this experiment, the subjects attended the cue (with eyes movement allowed) and fixed their gaze at the cross for 15-17 seconds and then the movie came on for 3 seconds. The movies were played on left and right monitors for 3 seconds before they return their gaze to the center screen to answer the questions like in the main experiment. After the movie, the subjects move their eyes back to the center monitor. The subjects completed 60 trials (3 runs of 20 trials each with break in between for subject comfort) each in this overt control task. Neon eye tracking glasses were used for all the control subjects. Eye gaze fixations typically stabilized within a second of eye movement initiation as in the main experiment (example eye traces from the main experiment are shown in figure 11).

Same processing pipeline was used as in the main experiment. However, we analyzed the run by setting two stimulus onset timing while modeling GLM: one with cue onset as stimulus onset timing and another with movie onset as the stimulus onset. GLM modeling was based on i) cue onset as stimulus onset timing (orienting phase of the experiment) and ii) movie onset timing (attending part of the experiment). This allowed us to independently model evoked responses to both the cue and the movie presentations. Each set of evoked responses were baseline subtracted with baseline (-2 to 0 seconds; pre stimulus) and concatenated temporally to show both sets of evoked responses together at a group level as shown in supplementary Fig. 1. A gap of uncertainty introduced by inter-trial interval jitter of 0-2 seconds is shown in gray color at 15-17th seconds.

*Supplementary results for IPL*

At cue onset, oxygenated and deoxygenated hemoglobin responses associated with the 'attend left' and 'attend right' conditions in IPL both show activation as well as clear divergence from each other, visually distinguishable at the group level (Supplementary Fig. 1). Notably, even as the subjects maintained eye fixation on their respective cued location (either −30° to the left or +30° to the right) for 15–17 seconds well after the completion of eye movement, the hemodynamic evoked responses remained sustained. At movie onset, the hemodynamic responses maintained their activation pattern (concatenated after baseline correction to capture relative changes) until movie offset. Following movie offset, when subjects moved their eyes back to the center screen at 0°, the evoked responses, reversed sign. Additionally, we observed a contralateral activation pattern in which the left angular gyrus was activated during the 'attend right' condition while the right angular gyrus was deactivated, and vice versa. These observations reveal that the IPL  encodes eye position and/or the locus of spatial attention, which are identical in the overt condition, and not eye movement.

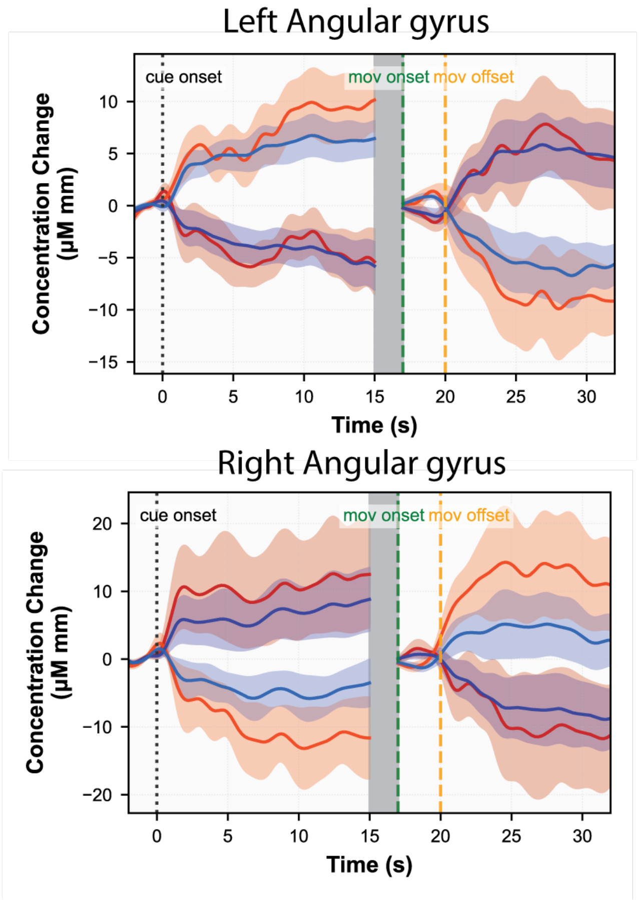


*Supplementary figure 1. ROI average Hemodynamic response functions on a group level (n=8) for the orient-wait-attend (control) experiment. The processing pipeline is the same as in figure 2 (please see supplementary methods section for more details).*

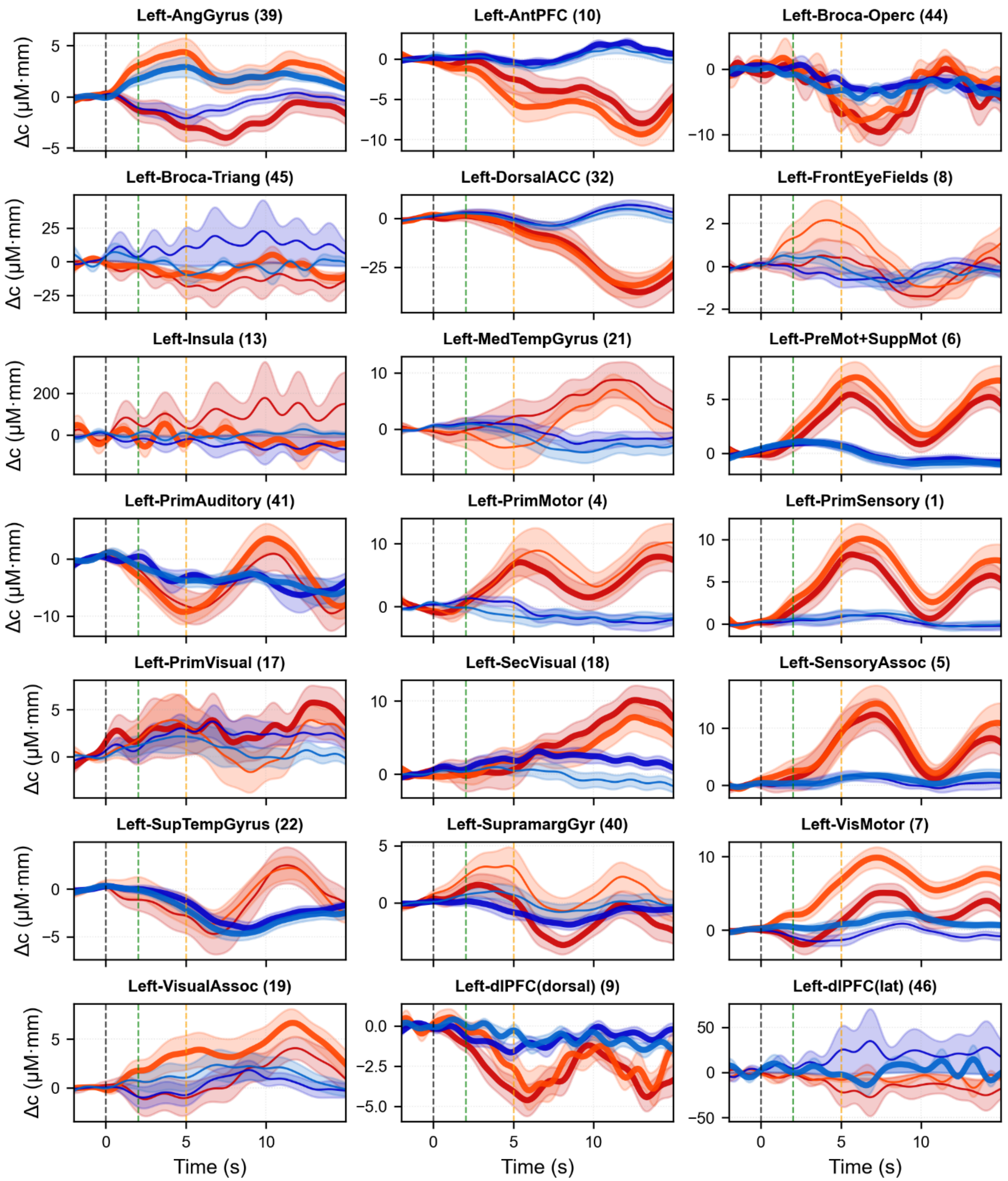


*Supplementary figure 2: ROI average Hemodynamic response functions on a group level. All ROIs from the left hemisphere in overt task are shown. Statistical significance was determined using t-test on the 2 seconds baseline (t=-2 to t=0) segment compared to the 2 seconds window starting cue (t=n to t=n+2, starting t=0) for both oxy and deoxy hemoglobin concentration. If any of the segments were statistically significant (p<0.05 after Bonferroni correction for multiple comparisons), the corresponding evoked response was marked as a statistically significant evoked response and indicated by bold lines.*

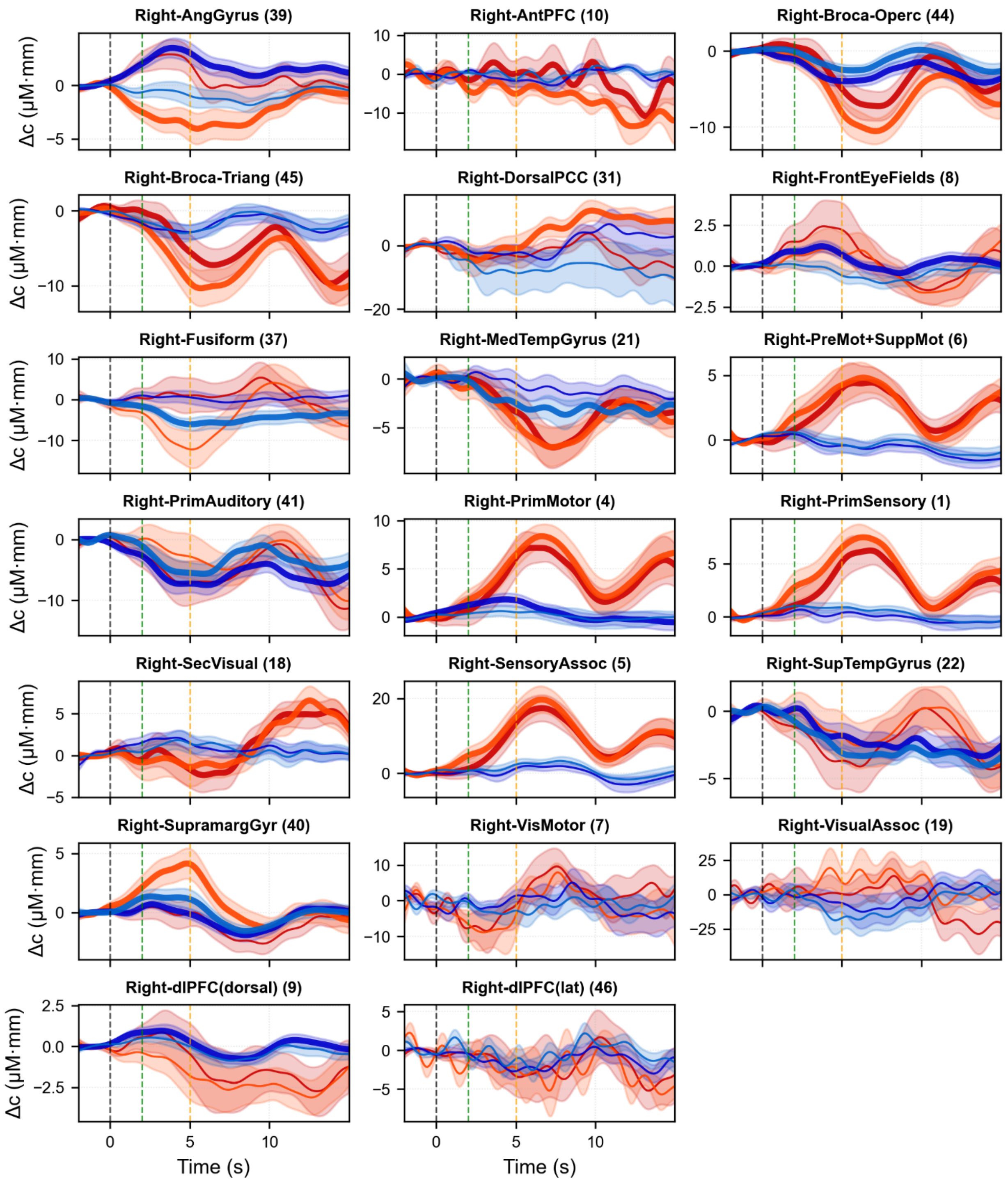


*Supplementary figure 3: ROI average Hemodynamic response functions on a group level. All ROIs from the right hemisphere in overt task are shown. Statistical significance was determined using t-test on the 2 seconds baseline (t=-2 to t=0) segment compared to the 2 seconds window starting cue (t=n to t=n+2, starting t=0) for both oxy and deoxy hemoglobin concentration. If any of the segments were statistically significant (p<0.05 after Bonferroni correction for multiple comparisons), the corresponding evoked response was marked as a statistically significant evoked response and indicated by bold lines.*

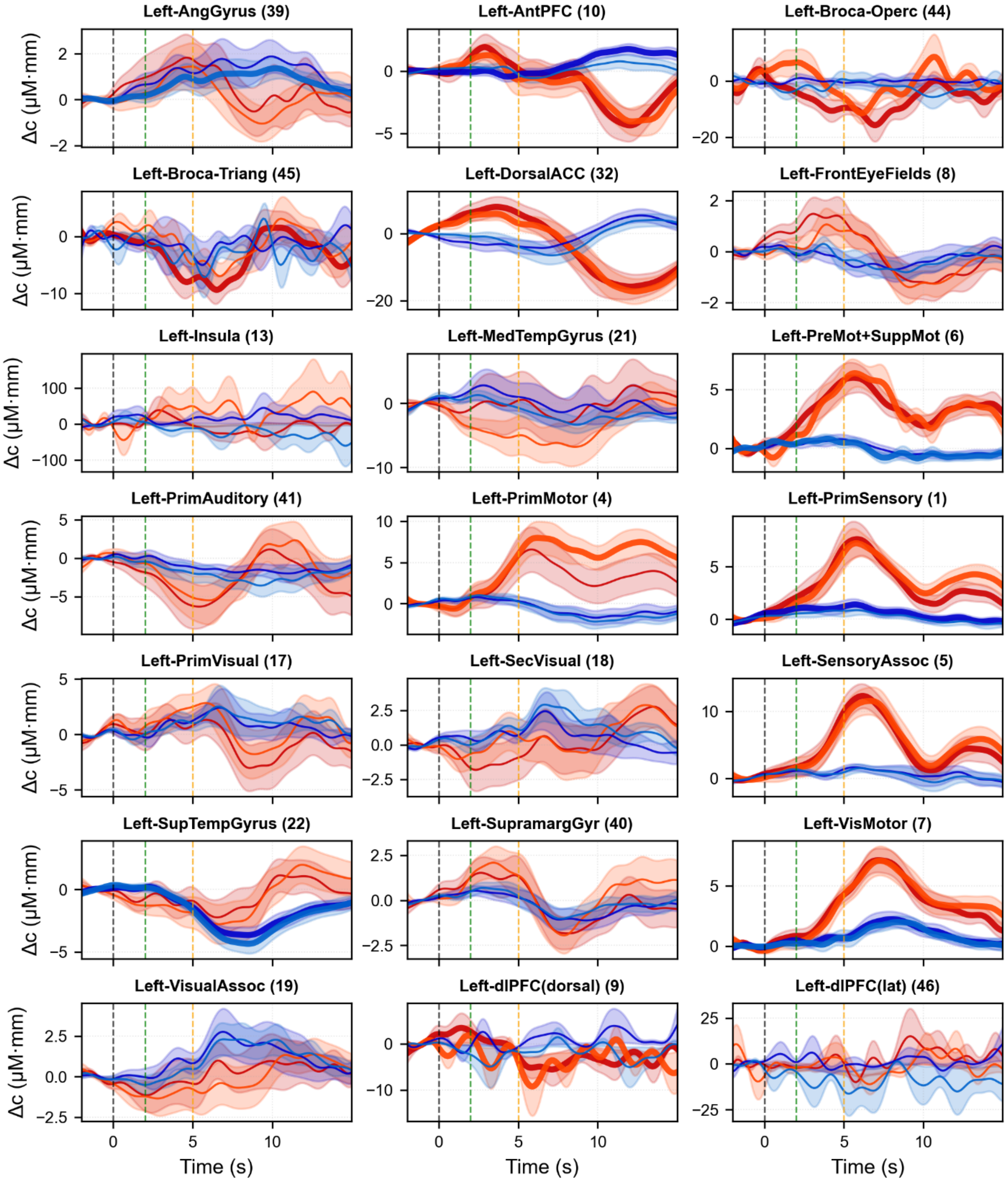


*Supplementary figure 4: ROI average Hemodynamic response functions on a group level. All ROIs from the left hemisphere in covert task are shown. Statistical significance was determined using t-test on the 2 seconds baseline (t=-2 to t=0) segment compared to the 2 seconds window starting cue (t=n to t=n+2, starting t=0) for both oxy and deoxy hemoglobin concentration. If any of the segments were statistically significant (p<0.05 after Bonferroni correction for multiple comparisons), the corresponding evoked response was marked as a statistically significant evoked response and indicated by bold lines.*

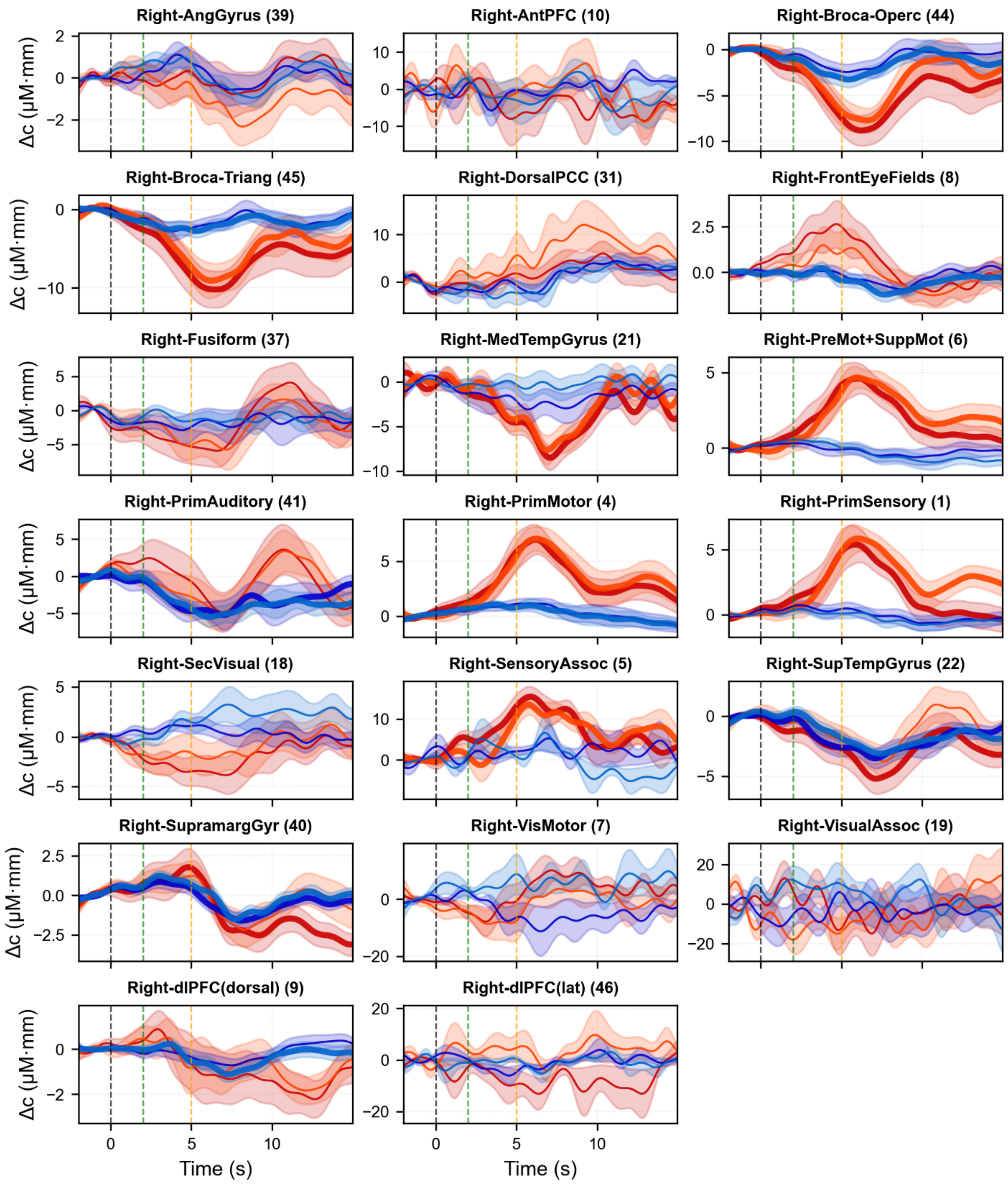


*Supplementary figure 5: ROI average Hemodynamic response functions on a group level. All ROIs from the right hemisphere in covert task are shown. Statistical significance was determined using t-test on the 2 seconds baseline (t=-2 to t=0) segment compared to the 2 seconds window starting cue (t=n to t=n+2, starting t=0) for both oxy and deoxy hemoglobin concentration. If any of the segments were statistically significant (p<0.05 after Bonferroni correction for multiple comparisons), the corresponding evoked response was marked as a statistically significant evoked response and indicated by bold lines.*

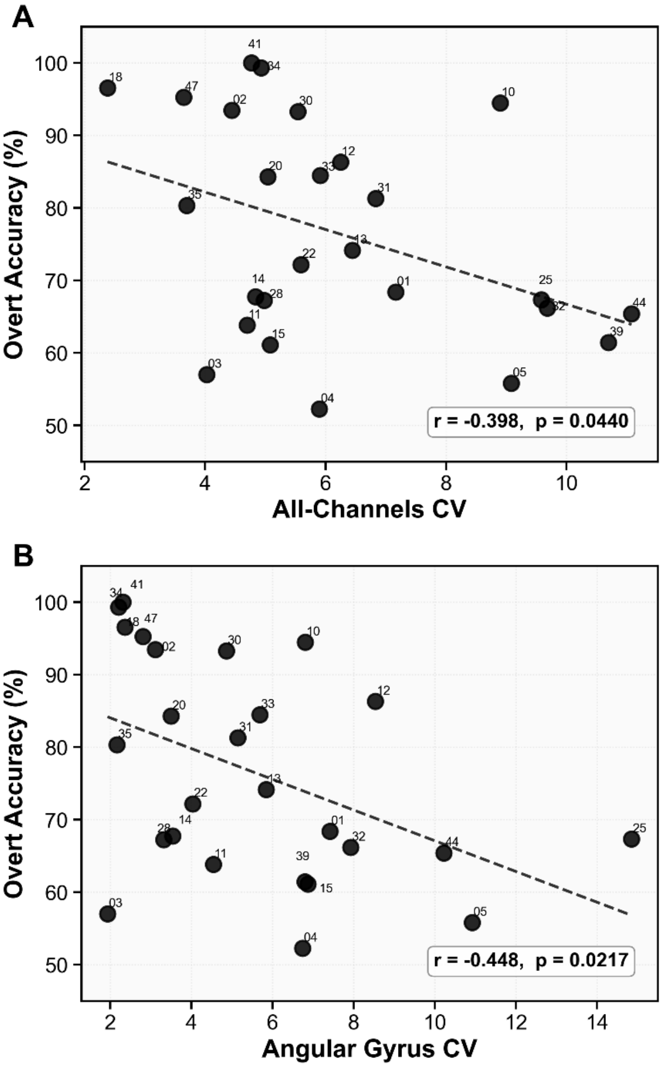


*Supplementary Figure 6. Relationship between within-subject variability of single-trial HbO responses and overt auditory attention classification performance. Within-subject variability was quantified per channel as the coefficient of variation (CV) = σ/|μ|, where μ and σ are the mean and standard deviation of the HbO response averaged over the 3–8 s post-stimulus window across all trials. Per-channel CVs were then averaged across (A) all recorded channels or (B) channels overlying the Angular Gyrus (Brodmann Area 39, left and right hemispheres pooled). Higher values indicate greater trial-to-trial variability relative to the mean response amplitude. Each dot represents one participant (N = 26); the y-axis shows overt classification accuracy (%). Dashed lines indicate the least-squares linear fit. Pearson r and p-values are shown in each panel.*

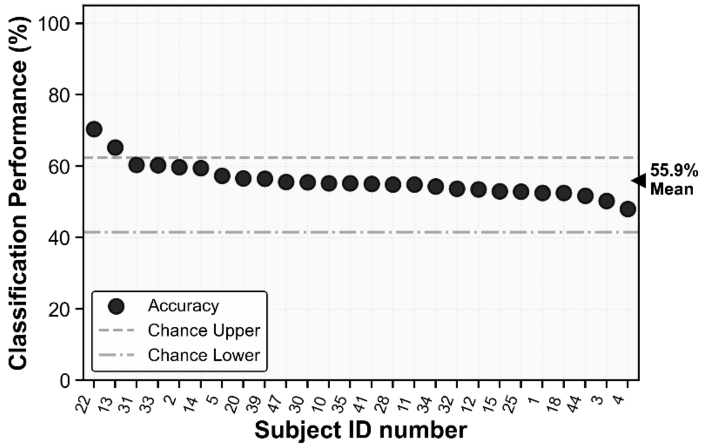


*Supplementary figure 7. Mean from 10 repetition of 5-fold cross validated classification accuracy (expressed in percentage) is shown for covert attention tasks. Two horizontal gray lines represent upper and lower confidence intervals for chance level after permutation test. All channels spanning the whole head coverage were considered for this portion of analysis.*

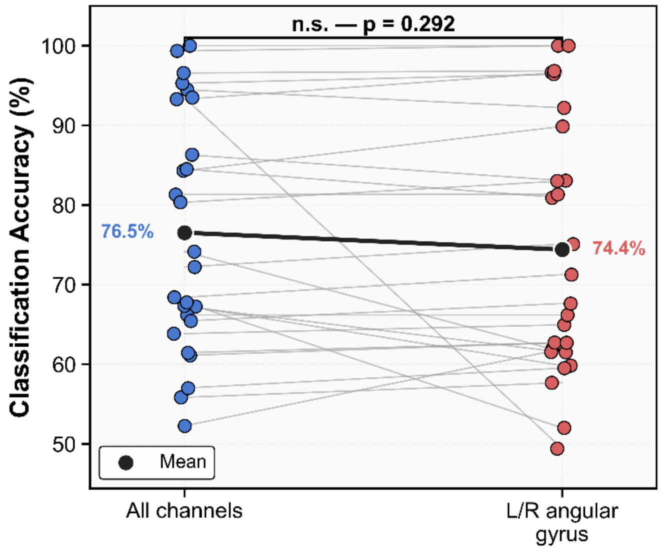

*Supplementary figure 8. Paired t-test among 26 pairs of all channels vs. left and right angular gyrus channels only. Classification accuracy does not significantly decrease when only left and right angular channels are considered.*

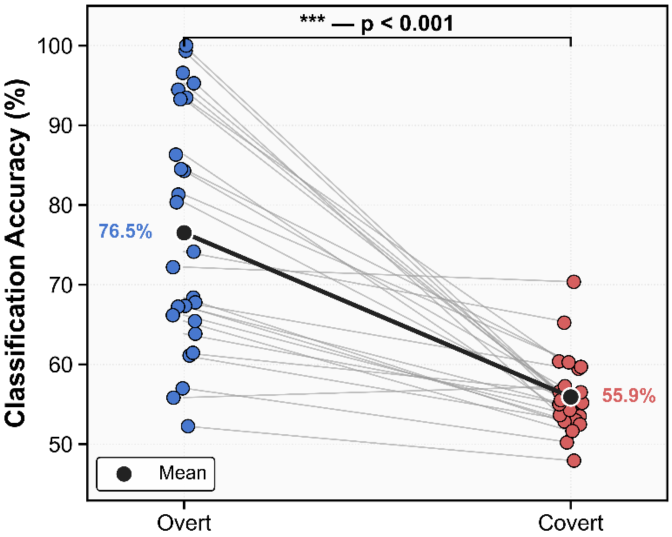


*Supplementary figure 9. Paired t-test among 26 pairs overt vs covert classification accuracy when all channels were considered. Classification accuracy significantly decreases in the covert task compared to the overt task.*
